# Topological Entropy Correlates with the Predictive Power of Multiplexed Ensemble Reservoir Computing

**DOI:** 10.64898/2026.02.04.703839

**Authors:** Suvankar Halder, Christopher M. Kim, Vipul Periwal

## Abstract

Modeling nonlinear, multiscale, and transiently chaotic biological processes remains a major challenge in computational biology. Traditional deep learning models, while powerful, require large datasets and lack mechanistic interpretability, limiting their effectiveness for time-resolved biological systems. Reservoir computing (RC) offers a promising alternative by leveraging the rich transient dynamics of fixed nonlinear systems, yet standard RC architectures struggle with high-dimensional biological data and complex temporal regimes. Here, we introduce Dynamical System Machine Learning (DynML), a multiplexed reservoir framework designed to model gene-expression dynamics in systems such as liver regeneration and *Drosophila* embryogenesis. DynML encodes biological signals using heterogeneous Lorenz reservoirs and employs a single global readout to capture stage-dependent dynamics with high predictive accuracy. We further show that reservoir topological entropy quantitatively predicts model performance, linking dynamical richness to biological forecasting accuracy. Beyond biological time-series modeling, we demonstrate the generality of DynML on the MNIST handwritten digit classification task using a Rössler-based chaotic reservoir, showing that fixed dynamical cores with linear readouts can also support high-dimensional static classification. Overall, DynML provides a scalable, interpretable, and computationally efficient framework that unifies biological time-series modeling and conventional machine-learning tasks within a single dynamical systems paradigm.

**Author summary:** Complex biological phenomena such as development, regeneration, and disease progression emerge from time-dependent gene-expression programs governed by nonlinear, multiscale dynamics. Capturing these dynamics remains challenging for conventional machine-learning approaches, which typically require large datasets and lack interpretability. In this study, we introduce Dynamical System Machine Learning (DynML), a modeling framework that leverages the transient dynamics of chaotic systems to learn and predict biological time series. DynML transforms gene-expression measurements into high-dimensional dynamical representations using ensembles of nonlinear reservoirs, enabling accurate prediction of future expression states with simple and interpretable linear readouts. We apply DynML to both synthetic dynamical systems and real biological datasets, including Drosophila embryonic development and human liver regeneration, where it achieves high predictive accuracy across multiple temporal transitions. Importantly, we show that the predictive performance of DynML is strongly linked to the topological entropy of the reservoir dynamics, providing a principled and quantitative measure of model expressiveness. Beyond biological time-series prediction, we demonstrate that the same dynamical framework can also classify static data, achieving strong performance on handwritten digit recognition. Together, our results establish DynML as a scalable and interpretable approach for modeling complex biological dynamics, and highlight how concepts from dynamical systems theory can guide the design of effective machine-learning models for biological data.

## 1 Introduction

Modeling complex biological systems remains a central challenge in systems biology and computational medicine. Biological processes involve intricate temporal dynamics, hierarchical regulation across molecular and cellular scales, and pronounced nonlinearity, which collectively hinder the effectiveness of traditional statistical or purely mechanistic modeling approaches [1, 2]. These systems often exhibit emergent behavior resulting from interactions among genes, signaling pathways, and environmental cues, requiring modeling frameworks that can accommodate high-dimensional, time-dependent, and noisy data [3, 4]. Recent advances in machine learning, particularly the integration of neural networks with dynamical systems theory, have enabled the development of hybrid models that are both predictive and biologically interpretable [5, 6].

Deep learning (DL) methods—including multilayer perceptrons (MLPs), convolutional neural networks (CNNs), recurrent networks such as Long Short-Term Memory (LSTM) and Gated Recurrent Unit (GRU), and more recently Transformer architectures—have been widely applied to biological data for tasks such as cell-type classification, gene regulatory inference, and phenotype prediction [7–11]. While these models excel at pattern recognition, they face key limitations when applied to biological dynamical systems. First, they require large labeled datasets, which are rarely available in time-resolved biological experiments. Second, deep models often lack mechanistic interpretability, making it difficult to connect learned representations to biological processes. Third, standard DL architectures struggle with long-term temporal dependencies, multiscale dynamics, and transient chaos—core features of developmental and regenerative systems [12]. These challenges motivate computational approaches that retain the expressive power of DL while better capturing the structure and dynamics of biological systems [11].

Reservoir computing (RC) is a computational paradigm rooted in recurrent neural networks and dynamical systems theory, particularly well-suited for processing time-dependent, nonlinear data. In this framework, the core component—known as the “reservoir”—consists of fixed, randomly connected recurrent units that project input signals into a high-dimensional dynamic space. Only the output weights are trained, significantly reducing computational complexity compared to fully trainable recurrent architectures [13–15]. By exploiting the reservoir’s rich temporal dynamics, this approach enables effective modeling of complex systems using simple linear readouts. Reservoir computing has been successfully applied in time-series forecasting, speech and pattern recognition, and classification tasks, owing to its ability to capture long-range dependencies and nonlinear transformations [16]. Furthermore, emerging research has explored hardware-based reservoirs—such as photonic and spintronic systems—offering promising directions for energy-efficient and high-speed computation [16]. Given these advantages, reservoir computing is increasingly recognized as a powerful tool for modeling biological phenomena characterized by temporal complexity, including gene regulation and tissue regeneration [17, 18]. A key feature of reservoir computing is the flexibility in selecting the reservoir substrate: instead of being limited to conventional neural networks, the reservoir can be implemented using a wide variety of nonlinear dynamical systems that exhibit rich transient dynamics and fading memory [19]. This design exploits the intrinsic computational capabilities of complex dynamical systems, where the reservoir itself performs temporal processing and only the readout layer needs to be trained. Reservoir computing frameworks inspired by dynamical systems have proven effective at capturing nonlinear temporal patterns. For instance, low-dimensional nonlinear systems such as a single time-delay system can utilize their intrinsic dynamics and memory to replace conventional recurrent neural network reservoirs [20], while a driven nonlinear pendulum can generate a rich temporal state space that serves as a computational reservoir [21]. The diversity of dynamical systems successfully employed as reservoirs demonstrates the universality of the reservoir computing paradigm, a point emphasized in recent comprehensive reviews that frame reservoir computing as a unifying framework across neural, physical, and dynamical substrates [22]. From quantum mechanical systems [23] to biological networks [24, 25], from optical fibers [26] to water waves [27], the common thread is the exploitation of rich, nonlinear dynamics with memory for computational purposes. This broad applicability suggests that reservoir computing represents a fundamental principle of computation in natural and artificial systems. The field continues to evolve with new physical substrates being explored, including DNA computing [28, 29], metamaterial structures [30, 31], and hybrid biological-electronic systems [32]. The key insight remains that computation is not limited to digital electronics but can emerge from any sufficiently complex dynamical system with appropriate input, dynamics, and readout mechanisms.

However, traditional RC approaches, such as echo state networks and liquid state machines, often face challenges when applied to complex biological systems. In these methods, input signals—such as gene-expression vectors—are projected into a high-dimensional reservoir, which is a recurrent dynamical system with fixed, randomly connected units. The reservoir transforms inputs into rich nonlinear trajectories that encode temporal patterns, and a single monolithic linear readout S is trained to predict the output at the next timepoint. While effective for general sequence modeling, standard RC has several limitations in biological contexts: it scales poorly with high-dimensional inputs and long sequences, struggles to capture transiently chaotic dynamics, and often lacks interpretability because the single readout combines contributions from all reservoir units without clear biological correspondence [11, 33].

To overcome these limitations, we introduce Dynamical System Machine Learning (DynML), a multiplexed reservoir architecture (see Fig. 1) designed for modeling nonlinear, transiently chaotic biological processes such as liver regeneration and *Drosophila* embryonic development. Here, multiplexed refers to the parallel coupling of multiple heterogeneous nonlinear dynamical systems into a single computational reservoir, rather than relying on a single recurrent network. Each subsystem evolves independently under the same input, and their collective states are concatenated to form a unified, high-dimensional representation of the system’s dynamics. In DynML, gene-expression measurements are fed into a parallel ensemble of heterogeneous Lorenz systems, generating rich dynamical trajectories that encode the system’s underlying state. The resulting multiplexed reservoirs collectively form a high-dimensional state vector, which is mapped to the next gene-expression state through a single global readout matrix S shared across all temporal transitions. By employing a single readout for all temporal transitions, DynML maintains interpretability while capturing the overall dynamics across developmental or regenerative stages. This formulation enables accurate prediction of both short-term fluctuations in *Drosophila* developmental waves and long-term regenerative trajectories in human liver tissue, while retaining the computational efficiency of fixed, heterogeneous reservoirs.

**Figure 1.**
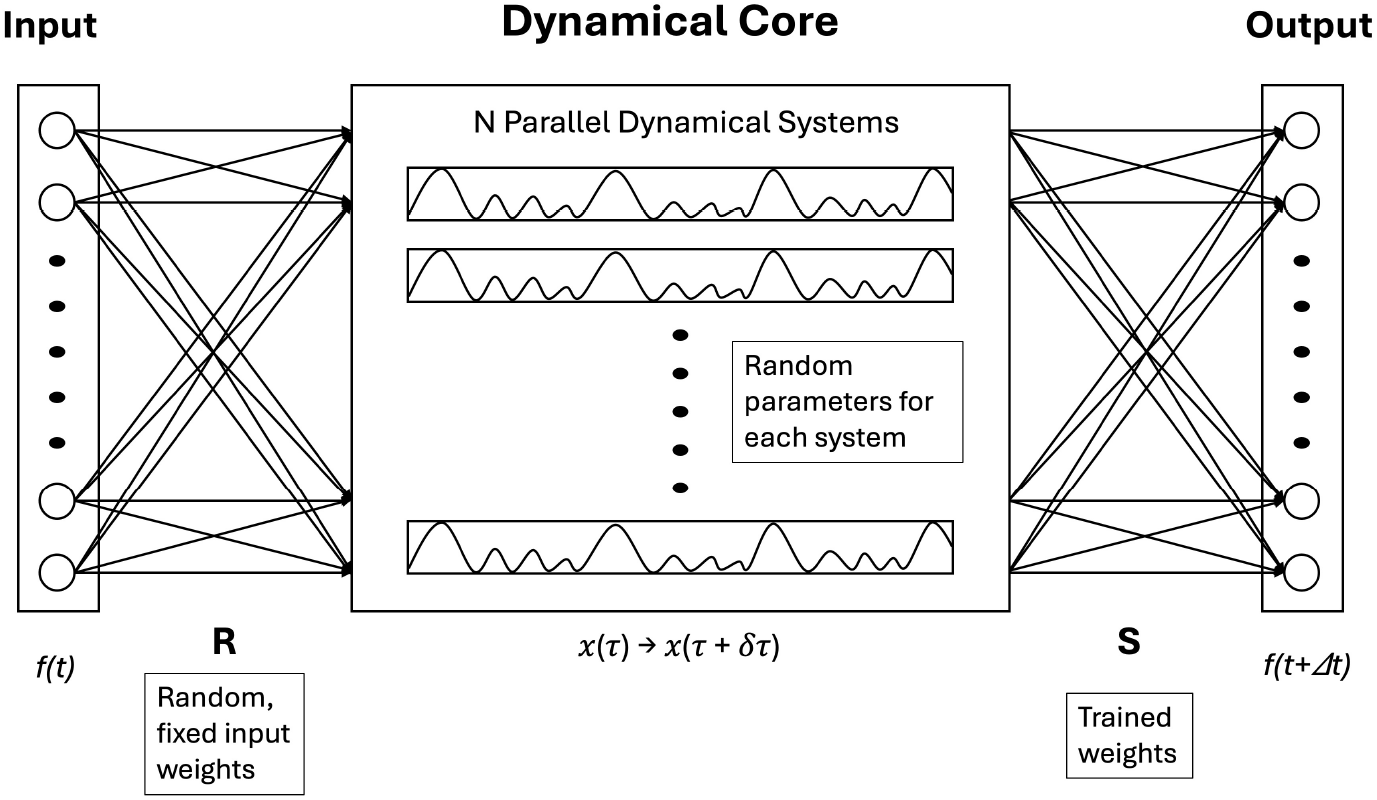
Architecture of the Dynamical System Machine Learning (DynML). The DynML consists of three layers: input, dynamical core (reservoir), and output. The dynamical core comprises multiple parallel dynamical systems with randomized parameters and initial conditions, forming a heterogeneous chaotic ensemble. The input signal is projected through a fixed random matrix *R* to perturb the initial states of each system. As the dynamical core evolves, its transient dynamics encode the input. Outputs are read at specific timepoints and linearly combined using a trained weight matrix *S*, enabling tasks such as prediction or classification by leveraging the system’s rich temporal behavior.

A central theoretical finding of this work is that topological entropy serves as a quantitative predictor of model performance. Reservoir ensembles operating in higher-entropy regimes generate a larger diversity of internal trajectories, providing a more expressive basis for mapping gene-expression patterns across time. Conversely, ensembles with low topological entropy exhibit diminished predictive accuracy, indicating insufficient dynamical complexity. We demonstrate that this relationship holds robustly across heterogeneous reservoirs and biological datasets, establishing topological entropy as a fundamental measure linking dynamical richness to predictive capability. Taken together, these contributions position DynML as a powerful and interpretable approach for modeling nonlinear biological time series. By uniting multiplexed reservoir computing with a principled dynamical-systems metric, the framework provides new insight into how biological complexity can be captured, predicted, and understood through the lens of nonlinear dynamics.

## 2 Methods

### 2.1 Mathematical Formulation of the DynML Architecture

We define DynML as a data-driven dynamical learning framework that maps an input vector **u** ∈ ℝ^*M*^ to a predicted output vector **ŷ** ∈ ℝ^*P*^, representing the system state at the next timepoint, using a heterogeneous ensemble of continuous-time chaotic systems as the dynamical core. In our implementation, this ensemble is constructed from multiple Lorenz systems.

#### 2.1.1 Input Projection

The input signal **u** is projected into the initial state of the dynamical core using a fixed, random linear transformation **R** ∈ ℝ^3*N* ×*M*^, where *N* is the number of dynamical systems:

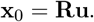

Here, **x**_0_ ∈ ℝ^3*N*^ represents the concatenated initial conditions [*x*_*i*_(0), *y*_*i*_(0), *z*_*i*_(0)] for all *i* = 1, …, *N* Lorenz systems.

#### 2.1.2 Reservoir Core Dynamics

Each dynamical core unit evolves according to a parameterized Lorenz system with parameters (*σ*_*i*_, *ρ*_*i*_, *β*_*i*_, *τ*_*i*_). The full core state **x**(*t*) ∈ ℝ^3*N*^ evolves in continuous time from the projected initial condition **x**_0_, governed by

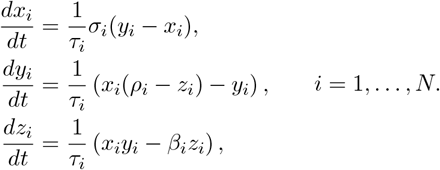

The Lorenz systems are numerically integrated over a fixed time horizon, and the final reservoir state **x**^∗^ ∈ ℝ^3*N*^ is used as the feature representation corresponding to the input vector **u**. Here, *τ*_*i*_ is a timescale parameter that controls the speed of evolution for each unit: smaller *τ*_*i*_ produces faster dynamics, while larger *τ*_*i*_ slows the evolution. By assigning different *τ*_*i*_ values across units, the dynamical core exhibits heterogeneous temporal behaviors, allowing the system to capture multiscale patterns in the input signals.

#### 2.1.3 Dynamical Core Readout

DynML employs a single linear readout matrix **S** ∈ ℝ^*P* ×3*N*^ that maps reservoir states to predicted outputs:

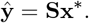

Importantly, the same readout matrix **S** is shared across all temporal transitions. During training, reservoir states corresponding to multiple temporal mappings (e.g., *t* → *t* + Δ*t* with variable Δ*t*) are pooled together, and a single global readout is learned to generalize across time.

This design enforces temporal consistency, improves statistical efficiency, and avoids overfitting to specific transition indices or time horizons.

#### 2.1.4 Training Objective

The readout matrix **S** is trained using linear least squares regression over all observed temporal transitions. Let **X** ∈ ℝ^3*N* ×*D*^ denote the matrix of reservoir states obtained from all training inputs across time, and let **Y** ∈ ℝ^*P* ×*D*^ denote the corresponding target outputs. The training objective is

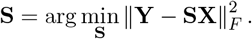

By learning a single readout across heterogeneous temporal transitions, DynML captures the underlying flow field of the dynamical system in state space while preserving interpretability and minimizing model complexity. This contrasts with transition-specific readouts, which require independent parameter estimation for each temporal mapping and are more susceptible to overfitting.

### 2.2 Topological Entropy of the Reservoir System

We quantified the complexity of the reservoir system by estimating its *topological entropy*, a measure of how unpredictable or chaotic the system dynamics are. Intuitively, topological entropy captures the rate at which initially nearby trajectories diverge over time: higher entropy indicates faster separation and more complex, less predictable behavior.

Formally, for a continuous map *f*: *X* → *X* on a compact metric space (*X, d*), the topological entropy *h*(*f*) can be defined in the sense of Bowen and Dinaburg as

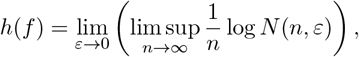

where *N* (*n, ε*) is the maximal cardinality of an (*n, ε*)-separated set with respect to the dynamical metric

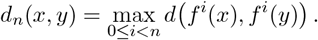

In simpler terms, *N* (*n, ε*) counts the largest number of trajectories of length *n* that remain distinguishable—i.e., separated by at least *ε*—over the entire time interval. This definition formalizes the idea that systems with higher entropy have a faster growth in the number of distinguishable trajectories and thus more complex dynamics.

Direct computation of *N* (*n, ε*) is generally infeasible for high-dimensional systems. Therefore, we employed a *finite-time, box-counting approximation* of the Bowen–Dinaburg entropy. Specifically, for each reservoir size *N*, the coupled Lorenz system was simulated with randomly drawn parameters (*σ, ρ, β*) and a time-scaling factor *τ*. The system was integrated using the Runge–Kutta (RK45) method at a finite number of time points. The trajectory of the 3*N*-dimensional state is denoted

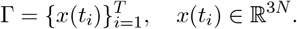

To approximate entropy, the trajectory was discretized at scale *ε* > 0 by partitioning the phase space into hypercubes of side length *ε*. Each point *x*(*t*_*i*_) was mapped to a symbolic state by

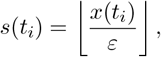

and the number of distinct symbolic states visited was denoted *N*_*n*_(*ε*). The finite-time entropy estimate was then computed as

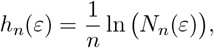

with *n* = *T* the trajectory length.

This procedure was repeated for multiple reservoir sizes *N* and multiple resolutions *ε*. For each configuration, entropy values were averaged across input samples. This approach provides a practical estimate of the system’s topological entropy and its dependence on reservoir size and phase-space resolution.

### 2.3 Chaotic Benchmark Systems

#### 2.3.1 Rössler System Dynamics

We employed the Rössler system as a low-dimensional continuous-time chaotic benchmark, as it is a canonical nonlinear dynamical system exhibiting a strange attractor and sustained chaotic dynamics. The system state (*x, y, z*) evolves according to

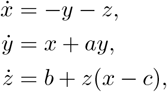

where *a, b*, and *c* are scalar parameters controlling the dynamical regime.

All simulations were performed in the chaotic regime using the standard parameter values *a* = 0.2, *b* = 0.2, and *c* = 5.7. A base initial condition (*x*_0_, *y*_0_, *z*_0_) = (0.1, 0.0, 0.0) was perturbed with additive Gaussian noise of standard deviation 10^−3^ to generate an ensemble of 2000 trajectories. The system was numerically integrated over a total time horizon of *t*_total_ = 120 using a high-precision Runge–Kutta solver (RK45) with tolerances rtol = 10^−9^ and atol = 10^−12^. To remove transient dynamics, the first 70 time units of each trajectory were discarded. Seven equidistant timepoints were then sampled between *t* = 70 and *t* = 120, yielding six temporal transitions for each state variable (*x, y, z*). The resulting dataset was organized such that rows correspond to state variables evaluated at successive timepoints (*x*_1_, …, *x*_7_, *y*_1_, …, *y*_7_, *z*_1_, …, *z*_7_), and columns correspond to distinct perturbed trajectories.

#### 2.3.2 Double Pendulum Dynamics

To benchmark DynML on a strongly chaotic mechanical system, we employed the planar double pendulum, a classical nonlinear system exhibiting extreme sensitivity to initial conditions. The state of the system is given by

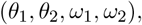

where *θ*_1_ and *θ*_2_ denote the angular positions of the first and second pendulum arms, and 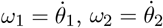 are the corresponding angular velocities.

The equations of motion are given by

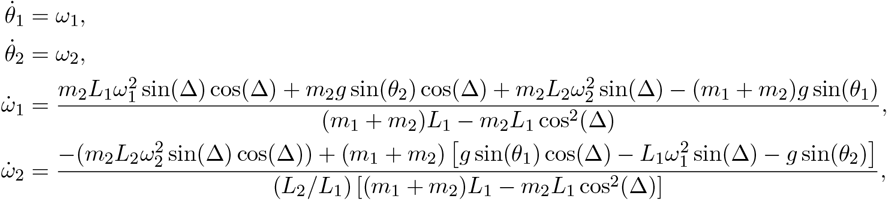

where Δ = *θ*_2_ − *θ*_1_, *m*_1_, *m*_2_ are the pendulum masses, *L*_1_, *L*_2_ are the arm lengths, and *g* is the gravitational acceleration.

All simulations were performed in a chaotic regime using parameters *m*_1_ = *m*_2_ = 1.0, *L*_1_ = *L*_2_ = 1.0, and *g* = 9.81. Trajectories were numerically integrated using a high-precision Runge– Kutta solver (RK45), and small perturbations to initial conditions were introduced to generate ensembles of chaotic trajectories. The base initial state (*θ*_1_, *θ*_2_, *ω*_1_, *ω*_2_) = (*π*/2, *π*/2 + 0.1, 0.0, 0.0) was perturbed using Gaussian noise with standard deviation 5 ×10^−3^ to generate 2000 trajectories. The system was integrated over *t*_total_ = 40 time units with solver tolerances rtol = 10^−9^ and atol = 10^−12^, discarding the first 10 units as transients. Seven equidistant timepoints were sampled, producing six transitions for each variable (*θ*_1_, *θ*_2_, *ω*_1_, *ω*_2_).

Both systems serve as benchmark dynamical substrates for evaluating DynML’s ability to learn and generalize chaotic temporal evolution from sparse, perturbed trajectory data using a multiplexed reservoir architecture. No explicit knowledge of the governing equations is provided to DynML during training; all predictions are learned purely from observed time-series data.

## 3 Results

### 3.1 Prediction of Chaotic Dynamics in Low-Dimensional Systems

We evaluated DynML on two canonical chaotic benchmark systems—the Rössler system and the planar double pendulum—chosen to represent low-dimensional continuous-time chaos and strongly nonlinear mechanical chaos, respectively. Details of the governing equations, simulation protocols, parameter regimes, and data organization are provided in the Methods section.

For both systems, DynML was trained exclusively on observed time-series data generated from ensembles of perturbed trajectories, without any explicit knowledge of the underlying equations of motion or system parameters. The task was to learn the temporal evolution of the system state and accurately predict future states across successive timepoints, despite sensitive dependence on initial conditions.

#### Data Preparation and Model Training

Input-output pairs were formed for each temporal transition *t* → *t* + 1, resulting in 18 tasks for the Rössler system and 24 tasks for the double pendulum. Each dataset was split into 80% training and 20% test samples and normalized using a standard scaler. DynML was configured with a multiplexed reservoir of heterogeneous Lorenz systems. For the Rössler dataset, *N* = 30 reservoirs were employed, while for the double pendulum, *N* = 200 reservoirs were used to accommodate the higher input dimensionality. Reservoir parameters (*σ, ρ, β*) were randomly sampled within narrow chaotic ranges: *σ* ∈ [9.9, 10.1], *ρ* ∈ [27.9, 28.1], *β* ∈ [2.65, 2.70], with a randomized time-scaling factor *τ*. Inputs were projected via a fixed random matrix, and reservoirs evolved for 30 time units to generate high-dimensional states. A single linear readout matrix was trained using ordinary least squares to map reservoir states to the target next timepoint.

#### Prediction Performance

DynML successfully learned the temporal evolution of both chaotic systems. For the low-dimensional Rössler system, predictions of (*x, y, z*) across six successive transitions captured the underlying attractor structure, with Pearson correlation coefficients near 1 and low mean squared errors on both training and test sets (top panel of Fig. 2). For the strongly chaotic double pendulum, DynML accurately forecasted the evolution of angular positions (*θ*_1_, *θ*_2_) and velocities (*ω*_1_, *ω*_2_) despite rapid divergence of nearby trajectories, achieving test-set correlations above 0.81 and statistically significant agreement with ground truth across all transitions (bottom panel of Fig. 2). These results demonstrate that DynML can robustly learn complex, nonlinear, and highly sensitive dynamics purely from sparse time-series observations.

**Figure 2.**
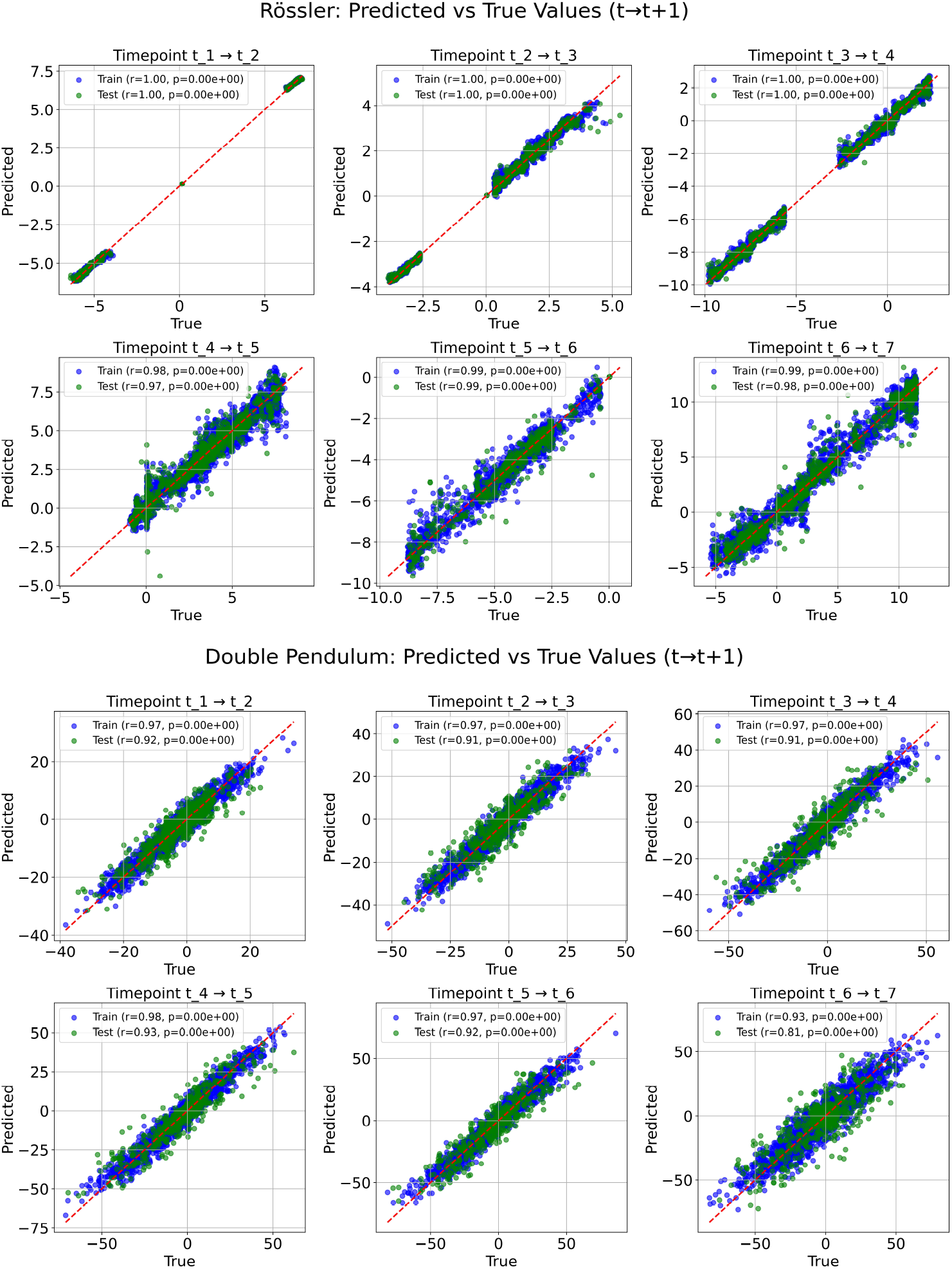
DynML predictions for chaotic systems. Top: Rössler system predictions. Scatter plots compare predicted versus true values for each temporal transition (*t* → *t* + 1) across all three variables (*x, y, z*). Blue points correspond to training samples, green to test samples. Pearson correlation coefficients and *p*-values are shown in the legends. **Bottom:** Double pendulum predictions. Scatter plots compare predicted versus true values for each temporal transition for the four state variables (*θ*_1_, *θ*_2_, *ω*_1_, *ω*_2_). Model captures highly nonlinear and chaotic dynamics across both training and test datasets, demonstrating DynML’s ability to generalize under strongly nonlinear regimes. Red dashed lines indicate the identity line.

### 3.2 Scaling DynML Performance with Reservoir Size

To examine how the size of the dynamical cores affects prediction performance and computational cost, we systematically varied the number of Lorenz subsystems *N* from 2 to 130 in increments of 4. For each value of *N*, a population of *N* Lorenz systems was instantiated with independently sampled parameters (*σ, ρ, β, τ*) drawn from narrow ranges around the canonical chaotic regime. A random input projection matrix *R* ∈ ℝ^3*N* ×*M*^ was generated for each *N* to map the normalized input states into the reservoir space. The computational complexity of DynML scales as 𝒪 (*N* ·*T*_int_) for reservoir evolution, where *N* is the number of Lorenz systems and *T*_int_ is the number of integration steps, plus 𝒪 (*D* · 3*N* · *P*) for readout training via least squares, where *D* is the number of training samples and *P* is output dimensionality.

DynML was evaluated on the double pendulum dataset using identical training and inference procedures across all experiments. Multiple random seeds were used to control the train–test split, reservoir parameter sampling, and input projection matrices. All reported results therefore represent the mean and standard deviation across seeds, ensuring robustness to stochastic initialization effects. For each input sample, the projected state was evolved through the multiplexed Lorenz reservoirs using numerical integration, and the final reservoir states formed a 3*N*-dimensional embedding. A linear readout trained via least-squares regression mapped this embedding to the target future state. Prediction accuracy was measured using the Pearson correlation coefficient, and computation time was recorded for each reservoir size *N*.

As shown in Figure 3, increasing the reservoir size consistently improves predictive accuracy for the chaotic system, with correlations rising rapidly for small*N* and gradually approaching saturation at larger reservoir sizes. In contrast, computation time increases overall with *N* but does not follow a strictly linear trend. Instead, it exhibits larger variability at smaller reservoir sizes, with substantial standard deviation, and gradually becomes more stable as *N* increases. The mean computation time displays a mildly sigmoidal growth, reflecting both initialization overhead at small *N* and increasing integration cost at larger *N*. Together, these results reveal a clear trade-off between predictive accuracy and computational efficiency and highlight an optimal operating regime in which high accuracy can be achieved while maintaining manageable and predictable computational cost.

**Figure 3.**
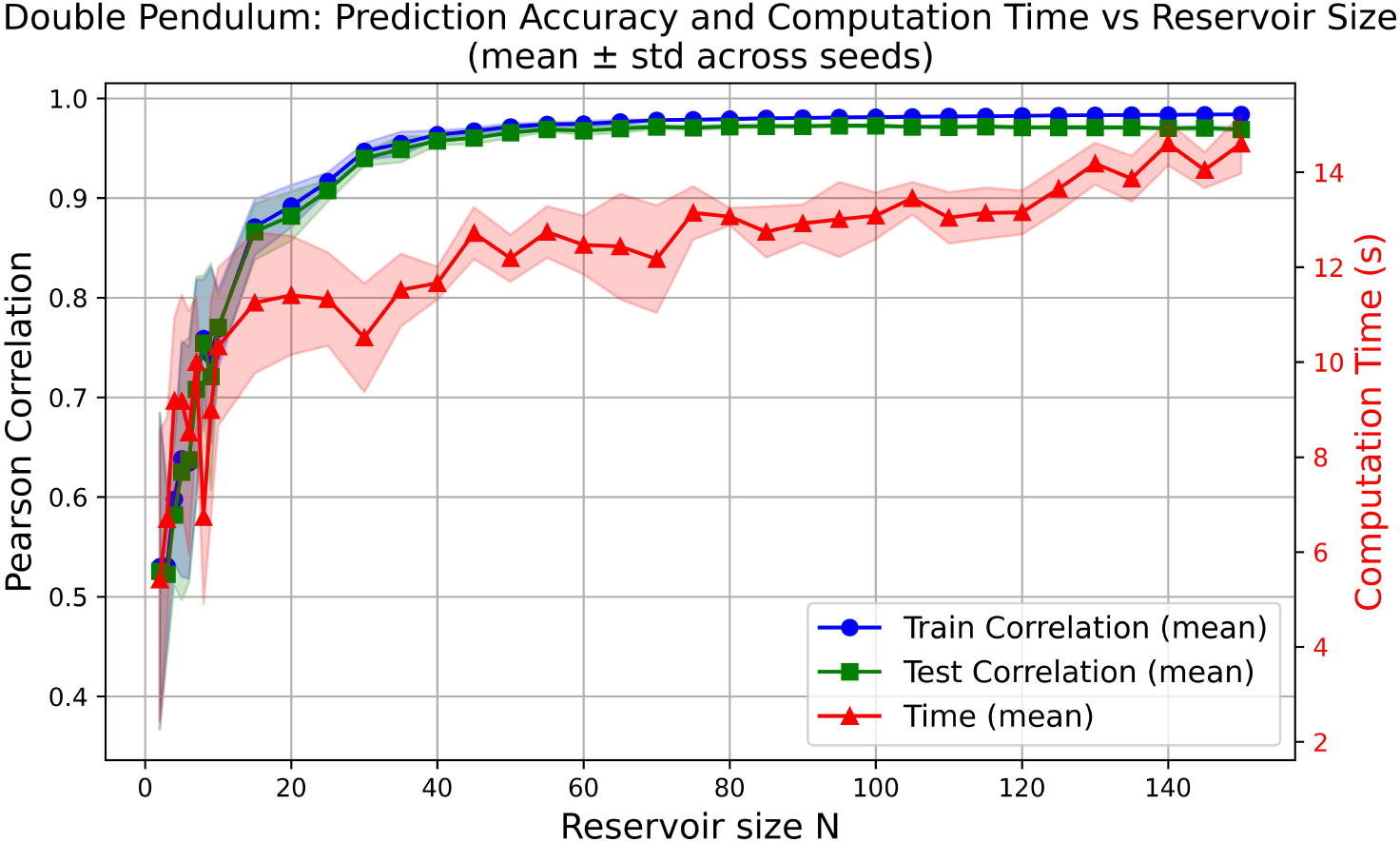
Double Pendulum: Prediction accuracy and computation time vs reservoir size. Mean Pearson correlation (train: blue, test: green) and mean computation time (red) are shown as functions of the number of Lorenz reservoir units (*N*), averaged across multiple random seeds. Shaded areas indicate one standard deviation across seeds. Increasing *N* improves predictive performance while increasing computation time, illustrating the accuracy–efficiency trade-off in DynML-based prediction of chaotic dynamics.

### 3.3 Prediction on Experimental Biological Data

Building on the highly encouraging results from chaotic-system benchmarks (Rössler system and double pendulum), we next applied DynML to real physiological datasets to evaluate its ability to model complex biological dynamics. Specifically, we tested the framework on *Drosophila* developmental gene-expression dynamics [34], and human liver regeneration time-series data [35, 36]. These datasets exhibit nonlinear, transient, and stage-specific behaviors that make them ideal for assessing DynML’s capacity to capture temporal structure and accurately predict biological state transitions.

#### 3.3.1 *Drosophila* Gene Expression Prediction Using DynML

We analyzed a publicly available spatiotemporal gene expression atlas of the *Drosophila* blastoderm [34]. The original VirtualEmbryo dataset contains quantitative mRNA expression measurements for 95 genes in 6078 individual nuclei across six embryonic timepoints, along with protein expression data for four gene products at selected stages. These high-resolution data enable systematic analysis of transcriptional dynamics during early embryonic development. After extracting spatial coordinates (*x, y,z*) and gene expression values, we filtered for genes with complete measurements across all six timepoints, resulting in a final set of 27 genes. Each gene contributes six expression values corresponding to successive developmental stages (*t*_1_, …, *t*_6_), yielding five temporal transitions (*t*_1_ → *t*_2_ through *t*_5_ → *t*_6_) per cell. The prediction task was to learn a mapping from gene expression at time *t* to expression at time *t* + 1, treating each cell–transition pair as an independent sample. To incorporate spatial context, the three-dimensional spatial coordinates of each cell were appended to the gene expression vector. Input features were standardized using training data only to prevent information leakage. The dataset was randomly split at the cell level into 80% training and 20% test sets, such that all six timepoints (and the corresponding five temporal transitions) from any given cell were assigned exclusively to either the training or the test set.

We employed a DynML framework with *N* = 150 randomly initialized Lorenz reservoirs. Each input vector (gene expressions plus spatial coordinates) was projected through a fixed random matrix *R* ∈ ℝ^3*N* ×*M*^ and injected as an initial condition into the reservoir. Although the prediction task is one-step (*t* → *t* + 1), the reservoir itself evolves according to its intrinsic continuous-time dynamics, producing a trajectory of high-dimensional states over the fixed integration period. These reservoir dynamics transform the static input into a rich temporal representation that captures nonlinear interactions between genes and spatial context. A linear readout was then trained on the training set to map the final reservoir state to the gene expression at the next timepoint, and evaluated on the held-out test set. In this way, the temporal dynamics of the reservoir provide a nonlinear feature transformation, enabling DynML to leverage temporal computation even for single-step predictions.

Model performance was evaluated using mean squared error and Pearson correlation between predicted and true expression levels. Figure 4 shows that both training (blue) and test (green) predictions cluster tightly around the identity line, indicating accurate prediction across all transitions. Test-set correlations exceed 0.94 for all transitions, with extremely low *p*-values, demonstrating strong statistical significance. These results confirm that DynML generalizes well to unseen cells and that the reservoir dynamics effectively encode the temporal and spatial structure of *Drosophila* gene expression.

**Figure 4.**
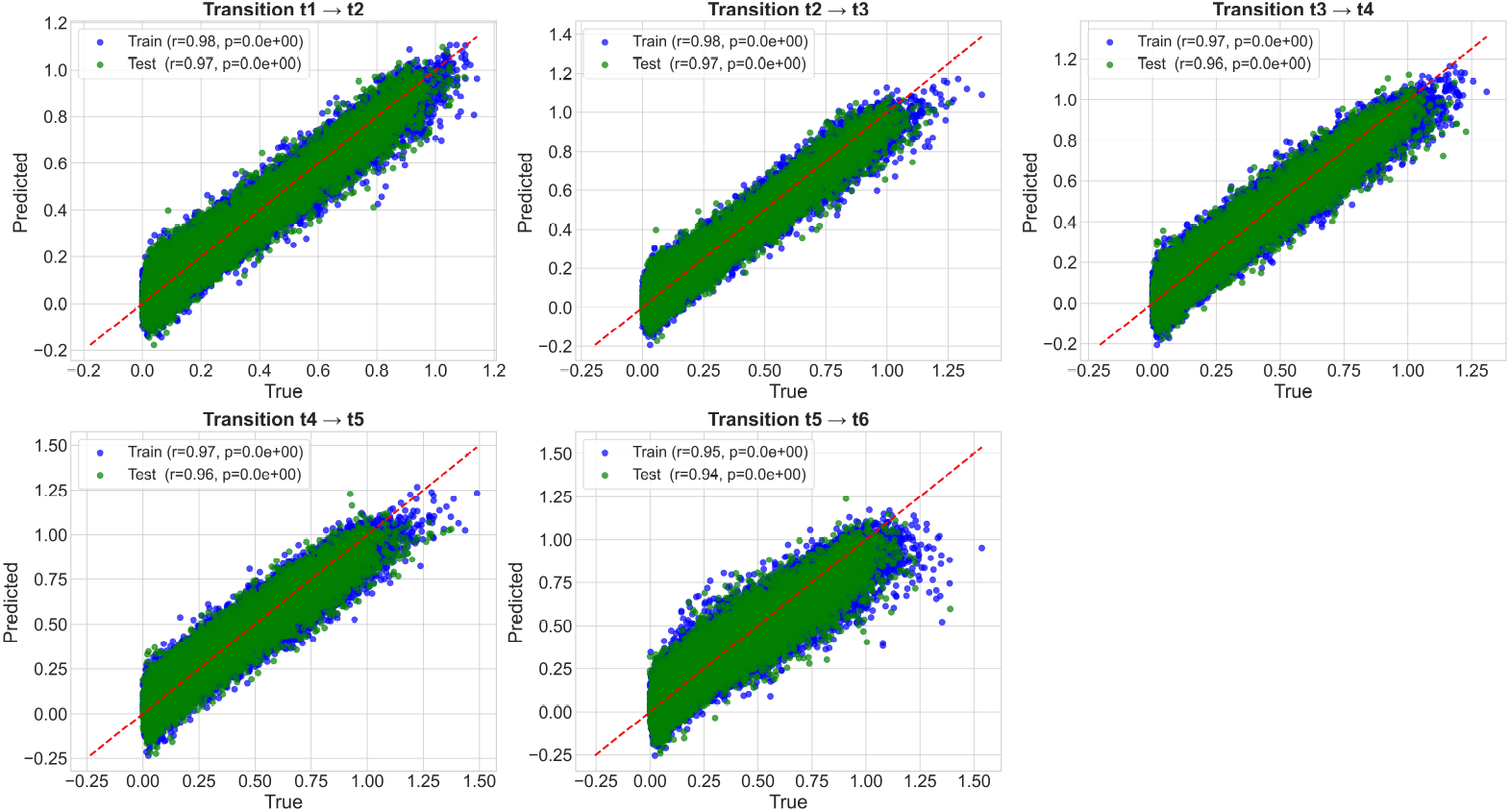
Prediction Performance Across Developmental Transitions in *Drosophila*. Scatter plots showing prediction performance for each temporal transition (*t* → *t*+1) across all 27 genes. Blue points: training set, green points: test set. Red dashed line: identity line. Pearson correlation coefficients (*r*) and *p*-values are displayed for each transition. Overall, DynML achieves strong predictive accuracy and generalization.

#### 3.3.2 Human Liver Regeneration Gene-Expression Prediction Using DynML

We evaluated DynML on a real physiological dataset describing human liver regeneration following partial hepatectomy. We used the transcriptomic data resource from Lawrence et al. [35], which provides longitudinal gene-expression measurements across 13 postoperative timepoints ranging from before surgery to one year after resection. The dataset captures regeneration-associated transcriptional programs across 12 patients, with genes grouped into biologically meaningful co-expression clusters [36]. These high-resolution temporal profiles make the dataset well suited for benchmarking predictive models of nonlinear, multi-phase regenerative dynamics.

For each patient, we extracted all available gene clusters (15 per patient) and randomly sampled one gene per cluster to construct a synthetic expression panel reflecting diverse regenerative programs. Repeating this process 6000 times produced a dataset of shape (6000, 15, 13), where each sample describes 15 cluster-representative genes tracked across 13 clinically defined regeneration stages. An 80/20 split was used to separate training and held-out test samples. We focused on two clinically relevant transitions that span distinct regenerative phases: 5 minutes → 1 day (acute inflammatory and early proliferative response) and 1 day → 3 months (mid-to-late remodeling phase). For each transition, the input vector consisted of gene expression at time *t*, and the output was the expression at time *t* + 1. After standardization, inputs were projected into a multiplexed reservoir of *N* = 150 heterogeneous Lorenz systems, and the reservoir dynamics were numerically integrated using an ODE solver. The evolving reservoir states produce a high-dimensional representation of the input, capturing nonlinear relationships between genes. A linear readout was then trained via least squares to map the reservoir states to gene expression at the next timepoint, and predictions were inverse-transformed back to the original units.

DynML achieved strong predictive performance across both transitions. Training and test predictions showed tight clustering around the identity line (Fig. 5), with high Pearson correlations for both transitions. For example, in a representative patient (Patient 1), the model achieved a test-set correlation of 0.87 for the 5 min → 1 day transition and 0.95 for the 1 day → 3 month transition, with training correlations consistently higher than those of the test sets; all correlations were highly significant (*p*-values near zero). Figure 5 shows results for four representative patients, while the corresponding results for the remaining patients are provided in the Supplementary Figure 1. These results indicate that DynML successfully captures the nonlinear temporal progression of liver regeneration and generalizes well to unseen samples. Taken together, this demonstrates the applicability of the reservoir-based framework to complex, multistage biological recovery processes.

**Figure 5.**
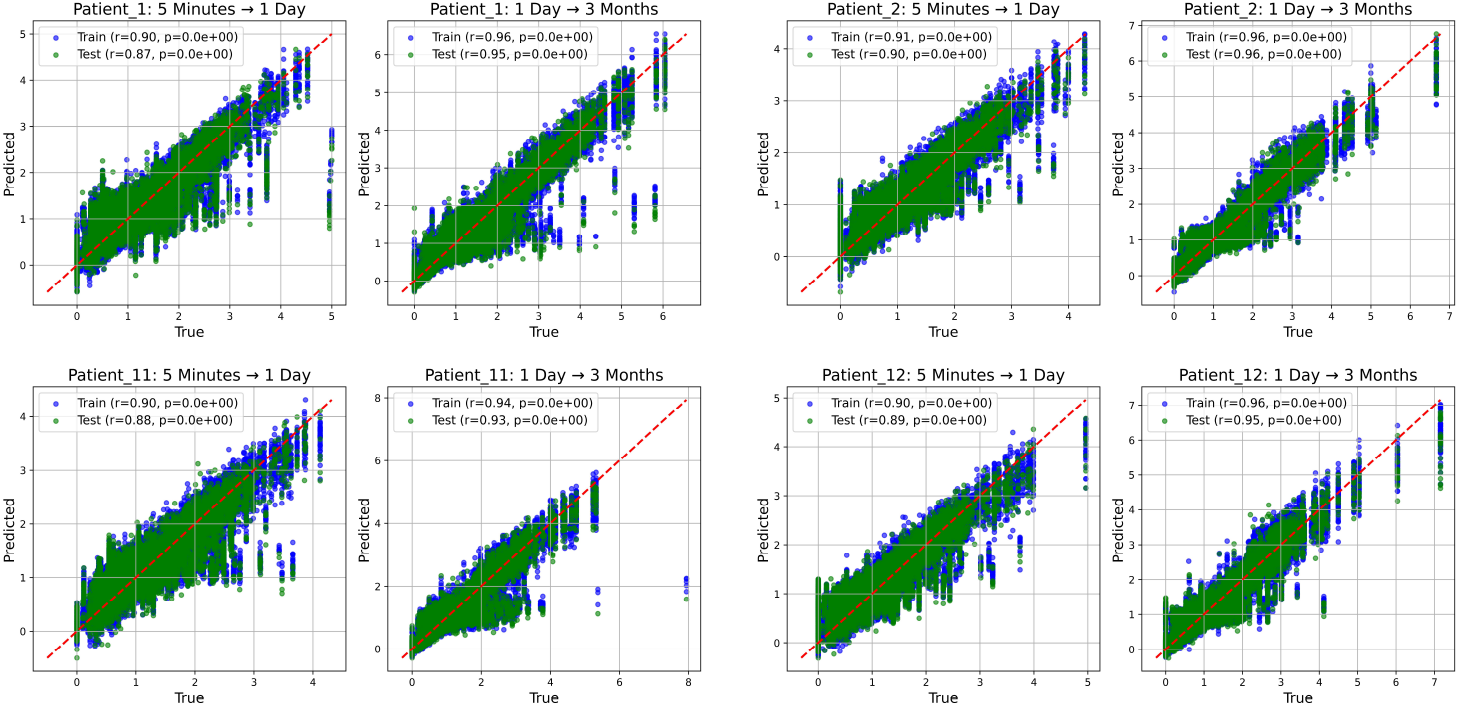
Prediction Performance Across Selected Temporal Transitions in Human Liver Regeneration. Scatter plots showing prediction performance for each temporal transition across all sampled genes for four patients. The task is to predict gene-expression levels at a later timepoint from the preceding timepoint. Blue points correspond to the training set and green points to the test set. For each transition, Pearson correlation coefficients (*r*) and associated *p*-values are displayed. The red dashed line denotes the identity line indicating perfect agreement between true and predicted values. Overall, the results demonstrate strong predictive accuracy and generalization of DynML across multiple patients and temporal transitions in liver regeneration.

### 3.4 Topological Entropy Increases with Reservoir Network Size

We quantified the system’s topological entropy to assess the dynamical richness and complexity of the reservoir networks. This metric provides a measure of how many distinct trajectories the system can generate, thereby reflecting the representational capacity of the network (for details see methods section).

We observed that the topological entropy increased monotonically as the number of reservoirs *N* was increased. For small *N*, the entropy values were low, indicating limited complexity in the state trajectories. As *N* grew, the entropy rose rapidly before reaching a plateau, suggesting that beyond a certain reservoir size, the addition of more reservoirs does not substantially increase dynamical complexity. The resolution parameter *ε* also influenced entropy estimates. At coarse resolutions (*ε* = 0.9), only large-scale trajectory features were captured, resulting in lower entropy. As *ε* decreased, finer distinctions between trajectories were resolved, leading to higher entropy estimates. For sufficiently small *ε*, entropy values stabilized, indicating that the effective complexity of the reservoir system had been fully captured. Overall, these results demonstrate that the reservoir system exhibits increasing dynamical richness with larger network size and that the topological entropy converges for sufficiently fine phase-space resolutions.

As shown in Figure 6, topological entropy estimates converge with respect to the box-counting resolution parameter *ε*. For all tested values of *ε* ranging from 0.9 to 10^−10^ (specifically *ε* ∈ {0.9, 0.5, 0.1, 0.05, 10^−2^, 10^−4^, 10^−6^, 10^−10^}), entropy increases monotonically with reservoir size *N* and saturates at large *N*. Critically, the relative ordering of entropy values across different *N* remains consistent regardless of *ε*, confirming that our entropy-based performance predictions are robust to numerical discretization. At finer resolutions (*ε* ≤ 0.1), entropy values stabilize, indicating that the effective dynamical complexity of the reservoir has been fully captured. This convergence across scales validates topological entropy as a reliable, resolution-independent measure of reservoir expressiveness.

**Figure 6.**
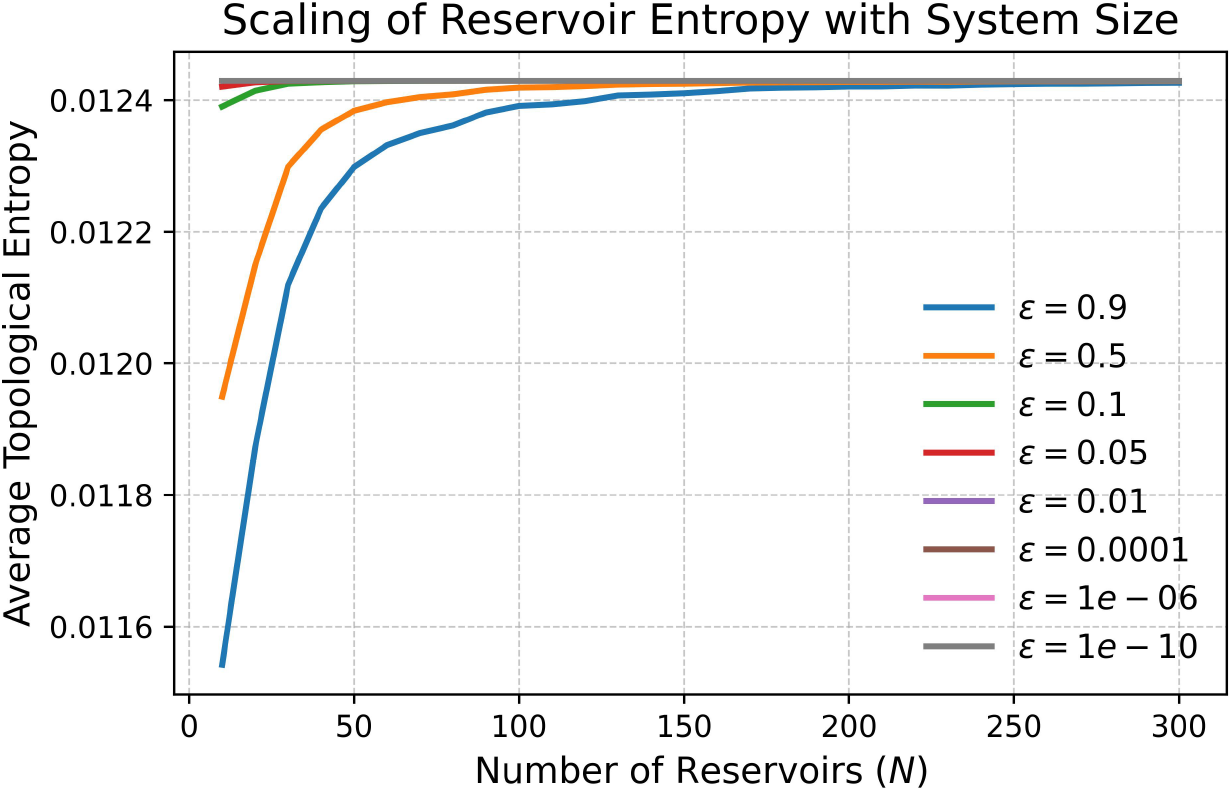
Topological entropy of the reservoir system. Average topological entropy as a function of the reservoir size *N* for different resolution scales *ε*. Entropy grows with *N* and saturates at larger reservoir sizes. As *ε* decreases, entropy increases but eventually stabilizes, reflecting convergence across scales.

### 3.5 Log-Scaled Reservoir Size Reveals Linear Entropy–Performance Relationship

To investigate how the internal dynamics of the Lorenz reservoir shape predictive performance, we quantified the dynamical richness of each reservoir configuration using topological entropy. Reservoir responses were simulated across varying sizes *N* (2–300) and multiple random seeds (*n* = 7) to ensure robustness. For each reservoir size, topological entropy was computed from the trajectories of all input samples and averaged across samples to obtain a single value characterizing that configuration.

Mean topological entropy and prediction accuracy (Pearson correlation) were then averaged across all seeds for each reservoir size. By plotting both quantities against log(*N*), we observed an approximately linear relationship: as the logarithm of the reservoir size increased, the mean entropy rose proportionally, and predictive performance improved in parallel. This trend was consistent across both the *Drosophila* developmental time-series dataset and the human liver regeneration dataset (Fig. 7). For the liver regeneration data, Fig. 7 shows results for a representative patient (Patient 3), while the corresponding analyses for the remaining 11 patients are provided in the Supplementary Figure 2.

**Figure 7.**
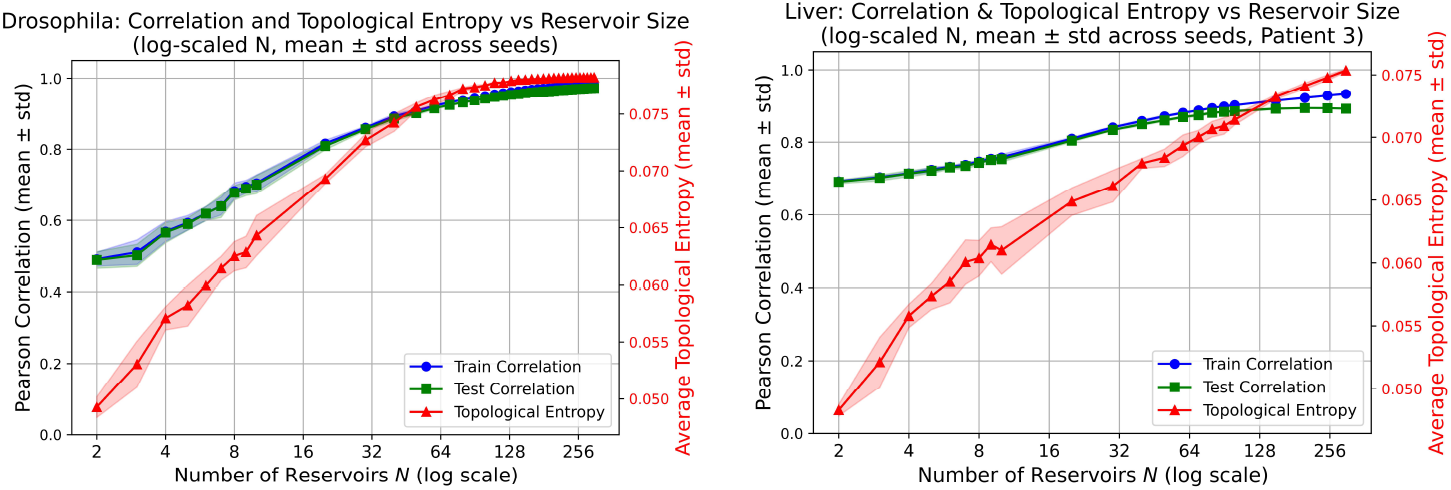
Linear relationship between log-reservoir size, topological entropy, and DynML performance. Mean topological entropy and predictive performance (Pearson correlation) were computed for each reservoir size *N* and averaged across seven independent random seeds. Across both the *Drosophila* and human liver regeneration datasets, mean train and test correlations increase approximately linearly with log(*N*), alongside the mean topological entropy of reservoir trajectories. Shaded regions indicate the standard deviation across random seeds. This reproducible trend establishes topological entropy as a robust criterion for selecting reservoir configurations and predicting DynML performance.

These results establish topological entropy as a quantitative and predictive indicator of reservoir expressiveness. Rather than relying on exhaustive hyperparameter searches, one can use the linear log-scale relationship to select reservoir sizes that maximize temporal prediction accuracy. The reproducibility of this trend across evolutionarily and mechanistically distinct biological systems underscores the generality of the approach and highlights the central role of dynamical complexity in modeling nonlinear gene-expression trajectories. The observed correlation between topological entropy and predictive accuracy can be understood through the lens of dynamical embedding theory. Higher entropy reservoirs generate a richer diversity of transient trajectories, effectively increasing the dimensionality and nonlinear expressiveness of the reservoir state space. This expanded representation allows the linear readout to separate complex input-output relationships that would be entangled in lower-entropy regimes. Mathematically, this suggests that topological entropy approximates the effective dimensionality of the reservoir’s functional basis, analogous to how kernel complexity determines separability in support vector machines.

### 3.6 Digit Classification with Rössler Reservoir-Based DynML

To demonstrate that DynML’s principles extend beyond temporal modeling, we conducted a proof-of-principle experiment on MNIST digit classification. This experiment serves not as a competitive benchmark against modern deep classifiers, but as validation that fixed chaotic cores with linear readouts can support high-dimensional static tasks when entropy is appropriately tuned. In contrast to earlier experiments that employed Lorenz reservoirs for modeling temporal biological dynamics, here we replaced the dynamical core with a highly nonlinear Rössler system to demonstrate that alternative chaotic systems also can provide expressive embeddings for static data.

A total of 20,000 images were used for training and 2,000 images for testing. Each input image was flattened and projected into a high-dimensional dynamical core consisting of *N* = 1500 parallel Rössler reservoirs with heterogeneous parameters. The resulting reservoir states formed nonlinear, high-dimensional representations of the input images, which were subsequently decoded using a linear readout implemented via logistic regression to predict digit labels. Despite the absence of task-specific feature engineering or deep architectures, DynML achieved a test accuracy of 87%, demonstrating that chaotic reservoir dynamics can effectively transform static spatial inputs into discriminative representations. These results confirm that the DynML framework is not restricted to temporal modeling and can successfully support high-dimensional static classification tasks.

Figure 8 illustrates representative test-set predictions, showing thirty randomly selected MNIST images along with their ground-truth and predicted labels. Misclassified digits are highlighted, providing qualitative insight into model performance. Together, these findings underscore the flexibility of DynML and highlight the central role of nonlinear dynamical cores in unifying temporal prediction and static classification within a single framework.

**Figure 8.**
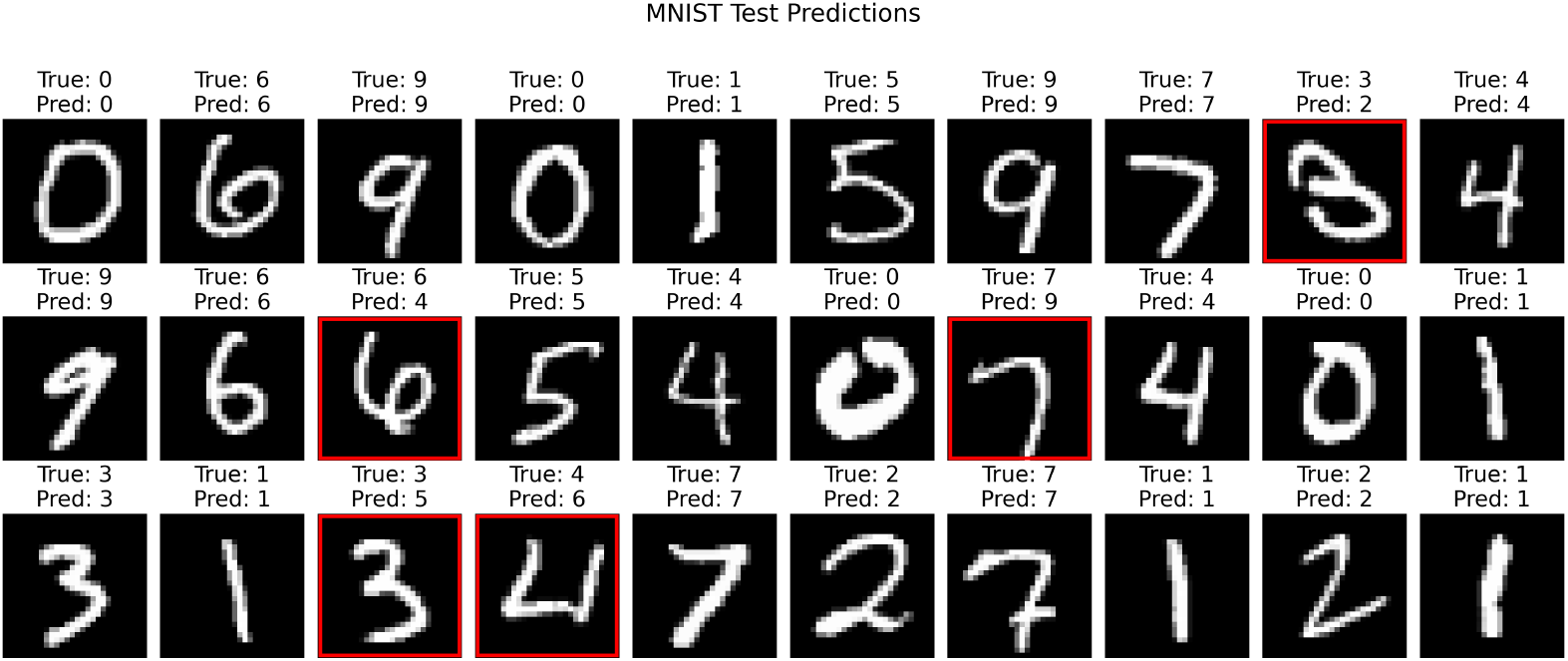
MNIST digit classification using DynML. Thirty randomly selected test images are shown with true labels and DynML-predicted labels. Images with incorrect predictions are outlined in red. Each input image is projected into a high-dimensional reservoir core (Rössler system, *N* = 1500), producing nonlinear embeddings that are linearly decoded to digit labels.

## 4 Discussion

In this study, we introduced DynML, a dynamical-systems-based machine learning framework for predicting and classifying complex temporal and spatial signals. DynML integrates ensembles of continuous-time chaotic systems as a heterogeneous reservoir core, enabling the capture of nonlinear dependencies and the generation of expressive high-dimensional representations. By employing multiplexed reservoirs with simple, task-specific linear readouts, the framework achieves strong predictive performance while remaining computationally efficient and interpretable.

DynML demonstrated robust performance across synthetic, biological, and classification tasks. For synthetic nonlinear dynamical systems, including the Rössler attractor and the double pendulum, the model accurately predicted state transitions from time *t* to *t*+1, capturing sensitive dependence on initial conditions and complex nonlinear dynamics. In biological applications, DynML successfully modeled temporal gene-expression transitions in *Drosophila* embryogenesis, achieving test-set Pearson correlations exceeding 0.94 across 27 genes, with highly significant statistical confidence. Similarly, in human liver regeneration data, DynML accurately captured patient-specific and time-resolved gene-expression dynamics, demonstrating strong generalization across individuals and temporal regimes. Beyond time-series modeling, DynML also performed high-dimensional static classification: when applied to the MNIST handwritten digit dataset, the framework transformed images into dynamical embeddings using a chaotic reservoir core and achieved 87% classification accuracy on held-out test data with a simple linear readout. Together, these results highlight the versatility of DynML across both temporal and spatial data domains.

The effectiveness of DynML arises from several key design principles. First, multiplexed chaotic reservoirs provide a rich and diverse set of nonlinear transformations that encode complex temporal and spatial structure more effectively than conventional feedforward or recurrent architectures. Second, the use of linear readouts enables efficient and stable training while preserving expressive power, allowing the model to map high-dimensional reservoir states to target outputs without extensive parameter optimization. Third, the modular architecture supports generalization across diverse problem domains, ranging from biological dynamics to image classification.

A central insight of this work is the role of intrinsic reservoir complexity in determining predictive performance. We showed that topological entropy provides a quantitative measure of reservoir expressiveness, with higher-entropy regimes consistently yielding improved prediction accuracy. Across both *Drosophila* development and human liver regeneration, reservoirs operating in higher-entropy regimes generated richer internal dynamics, enabling more accurate mappings between inputs and target states. In contrast, low-entropy reservoirs exhibited diminished performance, reflecting insufficient dynamical diversity. These findings establish topological entropy as a principled criterion for reservoir design and parameter selection, offering an alternative to exhaustive hyperparameter searches.

An important implication of these results concerns how reservoir expressiveness scales with computational cost. We observed that topological entropy increases approximately linearly with log(*N*), reflecting the multiplexed architecture of DynML, in which *N* heterogeneous but non-interacting chaotic subsystems operate in parallel. Increasing *N* therefore augments dynamical diversity by adding independent dynamical channels, rather than inducing a combinatorial expansion of the joint state space. As a consequence, dynamical complexity grows sublinearly with reservoir size, while the computational cost scales linearly in *N* due to independent low-dimensional simulations and the absence of backpropagation through time. This decoupling between expressiveness and optimization cost enables DynML to reach high-entropy, high-performance regimes using moderate reservoir sizes. Topological entropy thus serves not only as a predictor of predictive accuracy, but also as a practical guide for balancing model expressiveness against computational efficiency, clarifying how DynML achieves strong performance without requiring unnecessary increases in reservoir size.

While hierarchical [33] and multi-timescale [37] reservoir architectures have been explored, DynML differs fundamentally in three ways: (1) it uses continuous-time chaotic systems rather than discrete recurrent units, enabling direct exploitation of nonlinear flow dynamics; (2) it employs topological entropy as a predictive design criterion rather than a post-hoc performance metric; and (3) its multiplexed architecture with a single global readout maintains interpretability while scaling to biological dimensionality. This combination of continuous dynamics, entropy-guided design, and unified readout distinguishes DynML from prior heterogeneous reservoir approaches.

Despite these advances, several avenues remain for future development. Adaptive tuning of reservoir parameters or the integration of multiple chaotic system types may further enhance performance, particularly in noisy or highly nonlinear regimes [38]. Extending DynML to incorporate additional spatial or temporal context, such as multi-organ or single-cell datasets, could improve its applicability to complex biological systems. Moreover, incorporating uncertainty quantification into reservoir predictions would increase the framework’s utility in experimental and clinical settings.

Beyond developmental and regenerative biology, DynML has potential applications in physiological systems characterized by nonlinear and intermittently chaotic dynamics, such as cardiac electrophysiology. Cardiac rhythms emerge from multiscale interactions across molecular, cellular, and tissue levels, and pathological conditions such as arrhythmias are often associated with qualitative changes in underlying dynamical regimes. Chaotic reservoirs operating near critical entropy regimes may be particularly well-suited for detecting subtle deviations from normal cardiac dynamics. Our observation that topological entropy correlates strongly with predictive performance suggests that entropy-tuned reservoirs could amplify early irregularities in heartbeat time series, rendering incipient pathological signatures linearly separable at the readout level. Because DynML trains only the readout layer, such systems could adapt rapidly to patient-specific baselines while remaining sensitive to pathological deviations. Although not explored here, these considerations highlight a promising direction in which topological entropy serves not only as a predictor of model performance, but also as a control parameter for designing reservoir systems optimized for anomaly detection in complex physiological signals.

In summary, DynML provides a unified, interpretable, and computationally efficient framework for modeling, prediction, and classification of nonlinear, high-dimensional data. By coupling chaotic dynamical cores with simple linear readouts and principled measures of dynamical complexity, DynML bridges dynamical systems theory and machine learning, offering a powerful approach for understanding and forecasting complex biological and physical processes.

## Acknowledgments

This research was supported by the Intramural Research Program of the National Institute of Diabetes and Digestive and Kidney Diseases (NIDDK, ZIA DK075091-12) within the National Institutes of Health (NIH). The contributions of the NIH author(s) are considered Works of the United States Government. The findings and conclusions presented in this paper are those of the author(s) and do not necessarily reflect the views of the NIH or the U.S. Department of Health and Human Services.

## Materials & Correspondence

Correspondence and requests for materials should be addressed to Suvankar Halder.

## Data Availability and Code

All code required to replicate the analyses presented in this study is available at https://github.com/nihcompmed/DynML. The data used in this study are cited in the article.

## References

[1] H. Kitano, “Computational systems biology,” Nature, vol. 420, no. 6912, pp. 206–210, 2002.

[2] M. Sadria and V. Swaroop, “Discovering governing equations of biological systems through representation learning and sparse model discovery,” bioRxiv, pp. 2024–09, 2024.

[3] S. Huang, “Reprogramming cell fates: reconciling rarity with robustness,” Bioessays, vol. 31, no. 5, pp. 546–560, 2009.

[4] K. A. Janes and M. B. Yaffe, “Data-driven modelling of signal-transduction networks,” Nature reviews Molecular cell biology, vol. 7, no. 11, pp. 820–828, 2006.

[5] S. L. Brunton, B. R. Noack, and P. Koumoutsakos, “Machine learning for fluid mechanics,” Annual review of fluid mechanics, vol. 52, no. 1, pp. 477–508, 2020.

[6] C. Rackauckas, Y. Ma, J. Martensen, C. Warner, K. Zubov, R. Supekar, D. Skinner, A. Ramadhan, and A. Edelman, “Universal differential equations for scientific machine learning,” arXiv preprint arXiv:2001.04385, 2020.

[7] C. Angermueller, T. Parnamaa, L. Parts, and O. Stegle, “Deep learning for computational biology,” Molecular systems biology, vol. 12, no. 7, p. 878, 2016.

[8] J. Zou, M. Huss, A. Abid, P. Mohammadi, A. Torkamani, and A. Telenti, “A primer on deep learning in genomics,” Nature genetics, vol. 51, no. 1, pp. 12–18, 2019.

[9] X. Min, W. Zeng, S. Chen, N. Chen, T. Chen, and R. Jiang, “Predicting enhancers with deep convolutional neural networks,” BMC bioinformatics, vol. 18, no. Suppl 13, p. 478, 2017.

[10] C. Angermueller, H. J. Lee, W. Reik, and O. Stegle, “Deepcpg: accurate prediction of single-cell dna methylation states using deep learning,” Genome biology, vol. 18, no. 1, p. 67, 2017.

[11] X. Wang, P. Ma, J. Lian, J. Liu, and Y. Ma, “An echo state network based on enhanced intersecting cortical model for discrete chaotic system prediction,” Frontiers in Physics, vol. 13, p. 1636357, 2025.

[12] Y. Bengio, I. Goodfellow, A. Courville, et al., Deep learning, vol. 1. MIT press Cambridge, MA, USA, 2017.

[13] H. Jaeger and H. Haas, “Harnessing nonlinearity: Predicting chaotic systems and saving energy in wireless communication,” science, vol. 304, no. 5667, pp. 78–80, 2004.

[14] W. Maass, T. Natschlager, and H. Markram, “Real-time computing without stable states: A new framework for neural computation based on perturbations,” Neural computation, vol. 14, no. 11, pp. 2531–2560, 2002.

[15] A. Patharkar, F. Cai, F. Al-Hindawi, and T. Wu, “Predictive modeling of biomedical temporal data in healthcare applications: review and future directions,” Frontiers in Physiology, vol. 15, p. 1386760, 2024.

[16] G. Tanaka, T. Yamane, J. B. Héroux, R. Nakane, N. Kanazawa, S. Takeda, H. Numata, D. Nakano, and A. Hirose, “Recent advances in physical reservoir computing: A review,” Neural Networks, vol. 115, pp. 100–123, 2019.

[17] H. Jaeger, “The “echo state” approach to analysing and training recurrent neural networks-with an erratum note,” Bonn, Germany: German national research center for information technology gmd technical report, vol. 148, no. 34, p. 13, 2001.

[18] K. Nakajima and I. Fischer, Reservoir computing. Springer, 2021.

[19] K. Nakajima, “Physical reservoir computing—an introductory perspective,” Japanese Journal of Applied Physics, vol. 59, no. 6, p. 060501, 2020.

[20] L. Appeltant, M. C. Soriano, G. Van der Sande, J. Danckaert, S. Massar, J. Dambre, B. Schrauwen, C. R. Mirasso, and I. Fischer, “Information processing using a single dynamical node as complex system,” Nature communications, vol. 2, no. 1, p. 468, 2011.

[21] S. Mandal, S. Sinha, and M. D. Shrimali, “Machine-learning potential of a single pendulum,” Physical Review E, vol. 105, no. 5, p. 054203, 2022.

[22] M. Yan, C. Huang, P. Bienstman, P. Tino, W. Lin, and J. Sun, “Emerging opportunities and challenges for the future of reservoir computing,” Nature Communications, vol. 15, no. 1, p. 2056, 2024.

[23] K. Fujii and K. Nakajima, “Harnessing disordered-ensemble quantum dynamics for machine learning,” Physical Review Applied, vol. 8, no. 2, p. 024030, 2017.

[24] K. P. Dockendorf, I. Park, P. He, J. C. Príncipe, and T. B. DeMarse, “Liquid state machines and cultured cortical networks: The separation property,” Biosystems, vol. 95, no. 2, pp. 90–97, 2009.

[25] B. Jones, D. Stekel, J. Rowe, and C. Fernando, “Is there a liquid state machine in the bacterium escherichia coli?,” in 2007 IEEE Symposium on Artificial Life, pp. 187–191, Ieee, 2007.

[26] Q. Vinckier, F. Duport, A. Smerieri, K. Vandoorne, P. Bienstman, M. Haelterman, and S. Massar, “High-performance photonic reservoir computer based on a coherently driven passive cavity,” Optica, vol. 2, no. 5, pp. 438–446, 2015.

[27] C. Fernando and S. Sojakka, “Pattern recognition in a bucket,” in European conference on artificial life, pp. 588–597, Springer, 2003.

[28] D. Woods, D. Doty, C. Myhrvold, J. Hui, F. Zhou, P. Yin, and E. Winfree, “Diverse and robust molecular algorithms using reprogrammable dna self-assembly,” Nature, vol. 567, no. 7748, pp. 366–372, 2019.

[29] M. R. Lakin and D. Stefanovic, “Supervised learning in adaptive dna strand displacement networks,” ACS synthetic biology, vol. 5, no. 8, pp. 885–897, 2016.

[30] T. W. Hughes, I. A. Williamson, M. Minkov, and S. Fan, “Wave physics as an analog recurrent neural network,” Science advances, vol. 5, no. 12, p. eaay6946, 2019.

[31] B. G.-g. Chen, N. Upadhyaya, and V. Vitelli, “Nonlinear conduction via solitons in a topological mechanical insulator,” Proceedings of the National Academy of Sciences, vol. 111, no. 36, pp. 13004–13009, 2014.

[32] O. Feinerman, A. Rotem, and E. Moses, “Reliable neuronal logic devices from patterned hippocampal cultures,” Nature physics, vol. 4, no. 12, pp. 967–973, 2008.

[33] L. Manneschi, M. O. Ellis, G. Gigante, A. C. Lin, P. Del Giudice, and E. Vasilaki, “Exploiting multiple timescales in hierarchical echo state networks,” Frontiers in Applied Mathematics and Statistics, vol. 6, p. 616658, 2021.

[34] C. C. Fowlkes, C. L. L. Hendriks, S. V. Keränen, G. H. Weber, O. Rübel, M.-Y. Huang, S. Chatoor, A. H. DePace, L. Simirenko, C. Henriquez, et al., “A quantitative spatiotemporal atlas of gene expression in the drosophila blastoderm,” Cell, vol. 133, no. 2, pp. 364–374, 2008.

[35] M. C. Lawrence, C. M. Darden, S. Vasu, K. Kumano, J. Gu, X. Wang, J. Chan, Z. Xu, B. F. Lemoine, P. Nguyen, et al., “Profiling gene programs in the blood during liver regeneration in living liver donors,” Liver Transplantation, vol. 25, no. 10, pp. 1541–1560, 2019.

[36] S. Halder, M. C. Lawrence, G. Testa, and V. Periwal, “Donor-specific digital twin for living donor liver transplant recovery,” Biology Methods and Protocols, vol. 10, no. 1, p. bpaf037, 2025.

[37] G. Tanaka, T. Matsumori, H. Yoshida, and K. Aihara, “Reservoir computing with diverse timescales for prediction of multiscale dynamics,” Physical Review Research, vol. 4, no. 3, p. L032014, 2022.

[38] M. Lukoševičius and H. Jaeger, “Reservoir computing approaches to recurrent neural network training,” Computer science review, vol. 3, no. 3, pp. 127–149, 2009.

