## Supplementary materials for "Topological Entropy Correlates with the Predictive Power of Multiplexed Ensemble Reservoir Computing"

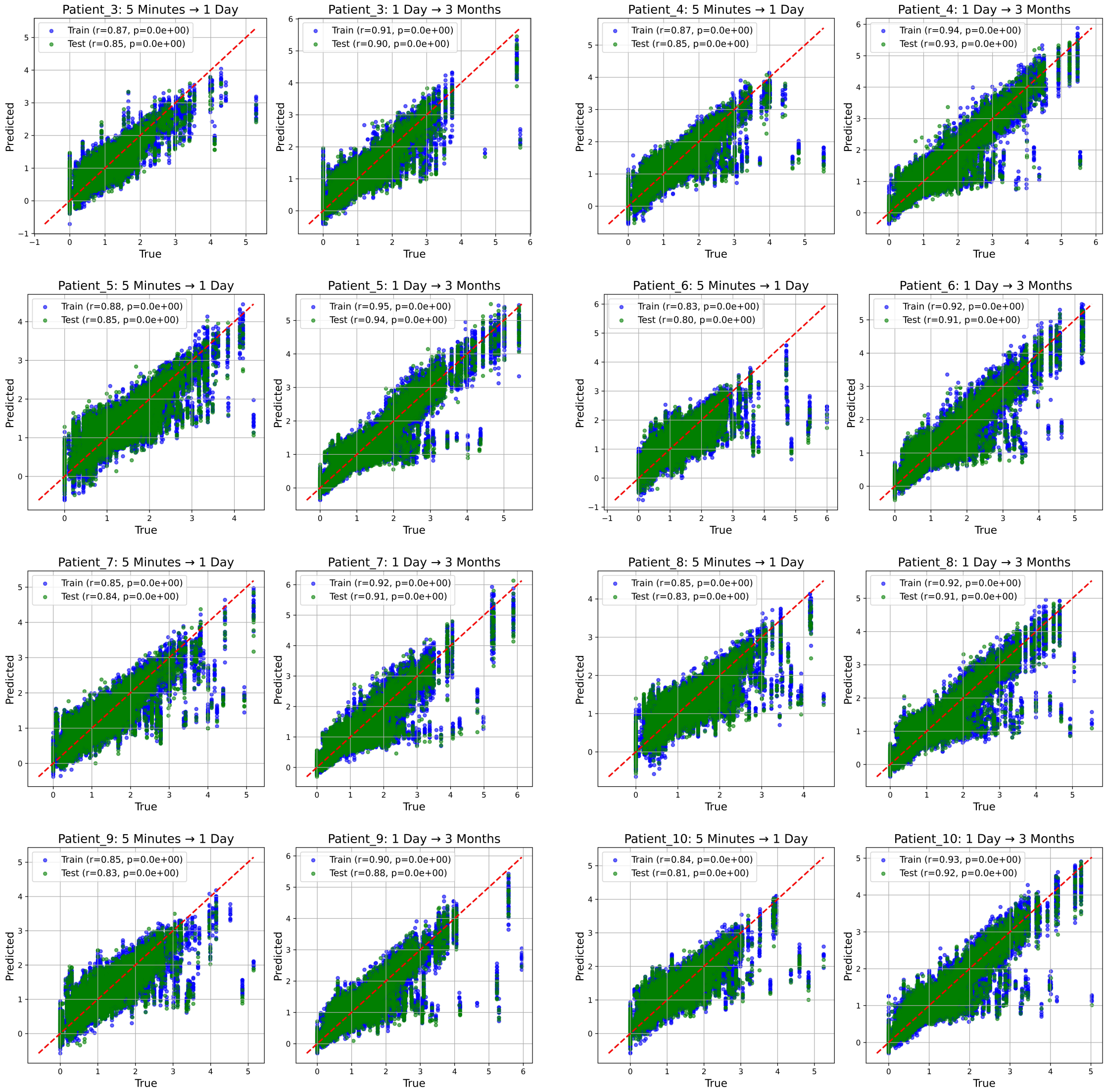


Figure 1: **Prediction Performance Across Selected Temporal Transitions in Human Liver Regeneration.** Scatter plots showing prediction performance for each temporal transition across all sampled genes for eight patients. The task is to predict gene-expression levels at a later timepoint from the preceding timepoint. Blue points correspond to the training set and green points to the test set. For each transition, Pearson correlation coefficients ($r$) and associated $p$-values are displayed. The red dashed line denotes the identity line indicating perfect agreement between true and predicted values. Overall, the results demonstrate strong predictive accuracy and generalization of DynML across multiple patients and temporal transitions in liver regeneration.

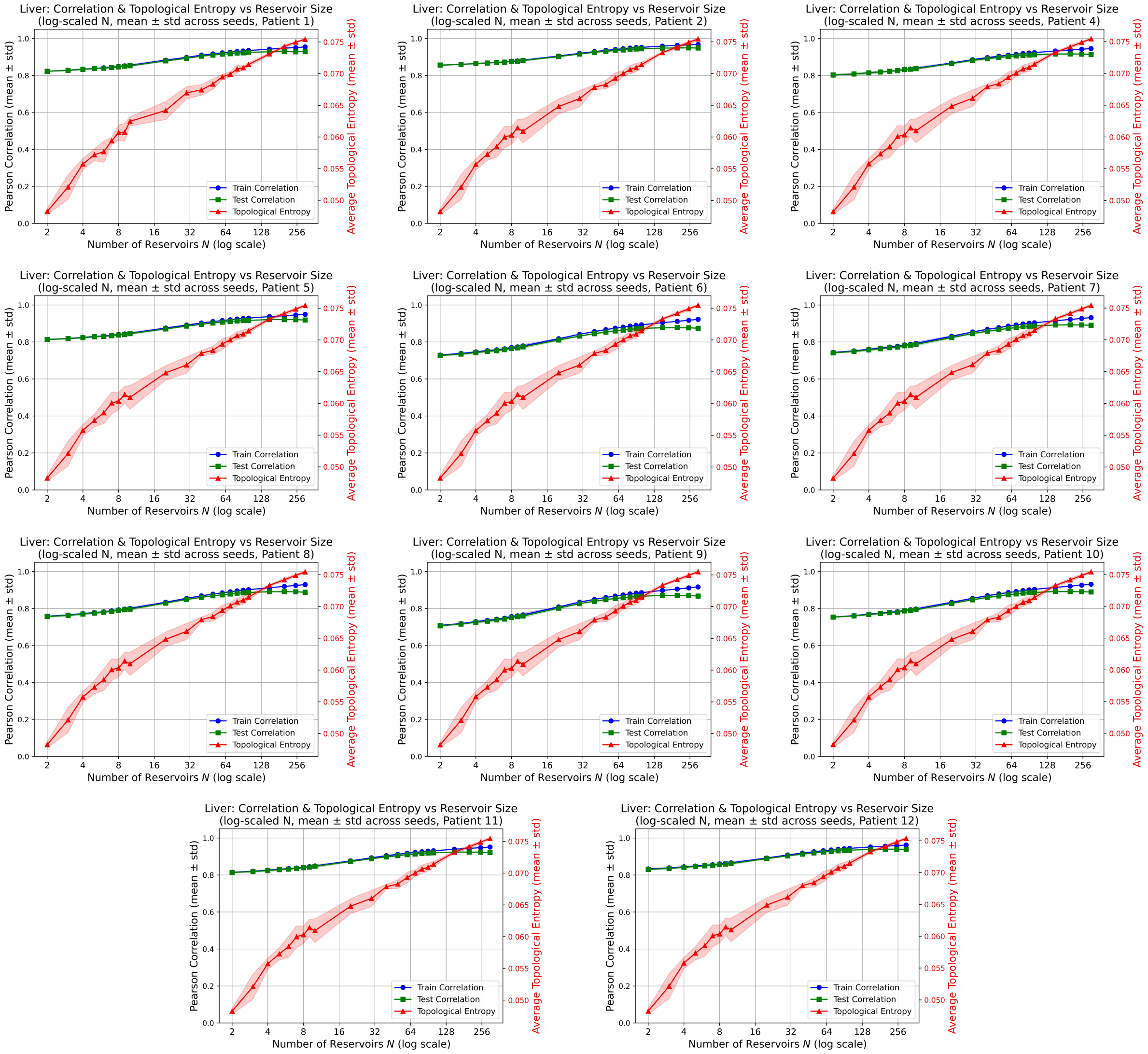


Figure 2: **Linear relationship between reservoir size, topological entropy, and DynML performance in human liver regeneration.** Mean topological entropy and predictive performance (Pearson correlation) were computed for each reservoir size $N$ and averaged across seven independent random seeds. For all patients shown, both train and test correlations increase approximately linearly with $\log(N)$, in parallel with increases in the mean topological entropy of reservoir trajectories. Shaded regions indicate the standard deviation across random seeds. This consistent trend across patients supports topological entropy as a robust criterion for selecting reservoir configurations and anticipating DynML performance.
